# Co-option of the transcription factor *SALL1* in mole ovotestis formation

**DOI:** 10.1101/2022.10.28.514220

**Authors:** Magdalena Schindler, Marco Osterwalder, Izabela Harabula, Lars Wittler, Athanasia C. Tzika, Dina Dechmann, Martin Vingron, Axel Visel, Stefan Haas, Francisca M. Real

**Affiliations:** Gene Regulation & Evolution. Max Planck Institute for Molecular Genetics, Berlin, Germany; Institute for Medical and Human Genetics, Charité - Universitätsmedizin Berlin, Berlin; Department for BioMedical Research (DBMR), University of Bern, Murtenstrasse 35, 3008 Bern, Switzerland; Department of Cardiology, Bern University Hospital, Bern, Switzerland; Environmental Genomics and Systems Biology Division, Lawrence Berkeley National Laboratory, 1 Cyclotron Road, Berkeley, California 94720, USA; Epigenetic Regulation and Chromatin Architecture. Max-Delbrück-Centrum für Molekulare Medizin (MDC) Berlin, Germany; Department of Developmental Genetics, Transgenic Unit. Max Planck Institute for Molecular Genetics, Berlin, Germany; Department of Genetics & Evolution. University of Geneva, Switzerland; Department of Migration and Immuno-Ecology, Max Planck Institute for Animal Behavior, Radolfzell, Germany; Department of Biology, University of Konstanz, Konstanz, Germany; Department of Computational Molecular Biology, Max Planck Institute for Molecular Genetics, Berlin, Germany; Department of Energy Joint Genome Institute, Berkeley, California 94720, USA; School of Natural Sciences, University of California, Merced, California 95343, USA

## Abstract

Changes in gene expression represent an important source for phenotypical innovation. Yet, how such changes emerge and impact the evolution of traits remains elusive. Here, we explore the molecular mechanisms associated with the development of masculinizing ovotestes in female moles. By performing comparative analyses of epigenetic and transcriptional data in mole and mouse, we identified *SALL1* as a co-opted gene for the formation of testicular tissue in mole ovotestes. Chromosome conformation capture analyses highlight a striking conservation of the 3D organization at the *SALL1* locus, but a prominent evolutionary turnover of enhancer elements. Interspecies reporter assays support the capability of mole-specific enhancers to activate transcription in urogenital tissues. Through overexpression experiments in transgenic mice, we further demonstrate the capability of *SALL1* to induce the ectopic gene expression programs that are a signature of mole ovotestes. Our results highlight the co-option of gene expression, through changes in enhancer activity, as a prominent mechanism for the evolution of traits.

## INTRODUCTION

Coordinated gene expression represents the cornerstone of developmental processes and homeostasis. In animals, transcription is mainly controlled by the action of cis-regulatory elements (CREs), such as enhancers, which control gene expression patterns with spatial and temporal precision. CREs control tissue-specific aspects of gene expression, acting in cooperation to constitute complex and pleiotropic gene expression patterns^1^. To exert their function, CREs are required to get into physical proximity with gene promoters, an operation mediated by the 3D folding of chromatin. CRE-promoter interactions are framed within topologically associating domains (TADs), large genomic regions with increased interaction frequencies, that are shielded from the regulatory influence of other genomic regions^2,3^.

Coding mutations usually modify the general function of a gene, thus inducing systemic effects that might be detrimental for the development of an organism. In contrast, mutations in CREs display tissue-specific effects, while preserving essential gene functions in other tissues. Consistently, the multiplicity of CREs can confer variations in expression patterns that contribute to gene pleiotropy, and support the rapid evolvability of these non-coding elements^4^. Variations in gene expression and function underlie the evolution of phenotypic traits and can be the basis for species adaptation. Indeed, mutations disrupting coding sequences have been associated with the emergence of certain traits, such as coat color or feathers^5^.

A prominent example of phenotypic evolution is observed in Talpid moles. In these species, female individuals consistently develop ovotestes, instead of ovaries^6^. These gonads are composed of ovarian tissue, supporting a fertile function, and a sterile testicular portion that secretes male hormones. These hormones exert a masculinizing effect in female moles, increasing muscle strength and aggression, in line with their adaptation to subterranean environments. In a previous study, we demonstrated that the evolution of ovotestes is associated with the reorganization of TADs, which alter CRE-promoter interactions and gene expression patterns^7^. In particular, a large inversion relocates active enhancers in the vicinity of the pro-testicular gene *FGF9*, whose ectopic expression in female gonads leads to meiosis inhibition and masculinization. In addition, a duplication of enhancer elements is associated with the increased expression of *CYP17A1* encoding an enzyme for male hormone synthesis and increases muscle strength. While the observed regulatory changes at these loci explain partially the mole phenotype, it is plausible that additional mechanisms contribute to the evolution of this trait.

In this study, we further investigate the molecular mechanisms associated with mole ovotestis development. Using comparative epigenetic and transcriptional approaches in mole and mouse, we identify the transcription factor *SALL1* as an early marker of mole ovotestis development. We observe that *SALL1* has been co-opted in the formation of XX testicular tissue in the Iberian mole *Talpa occidentalis*. Our finding is further supported by expression analyses in closely related species developing normal ovaries, like shrews and hedgehogs. We determine the regulatory landscape of this gene, highlighting an evolutionary conserved TAD structure that undergoes prominent variation at the internal enhancer composition. Through *in vivo* interspecies reporter assays, we highlight the potential of enhancer variation to evolve new activity domains in moles. By using transgenic mice that overexpress *Sall1* in ovaries, we demonstrate the capacity of this factor to activate gene expression programs that are distinctive of mole ovotestis formation. Altogether, our results shed light on the molecular basis of a unique trait, putting forward the important role of regulatory variation in evolution.

## RESULTS

### Evolutionary conservation of mammalian gonadal enhancers

CREs represent a major source of tissue-specific gene expression. We previously explored the regulatory landscape of mole developing gonads, at an early postnatal stage (7 days *post-partum* - stage P7). At this developmental time-point, testicular and ovarian tissues from female ovotestes are first morphologically discernable and can be clearly microdissected (**Figure 1A**). Furthermore, Leydig cells of the testicular part differentiate and produce testosterone, whereas meiosis initiates in the ovarian part, an event considered as one of the earliest signs of female gonadogenesis in mammals^8^. We identified regulatory elements in mole gonads by performing ChIP-seq experiments against a combination of histone marks, H3K27Ac together with H3K4me1 and H3K4me3, for the distinction of enhancers and promoters, respectively. By using the tool CRUP^9^, we combined these datasets in each sampled tissue to call and rank active regulatory regions according to their enhancer probability score (**Figure 1B**).

**Figure 1.**
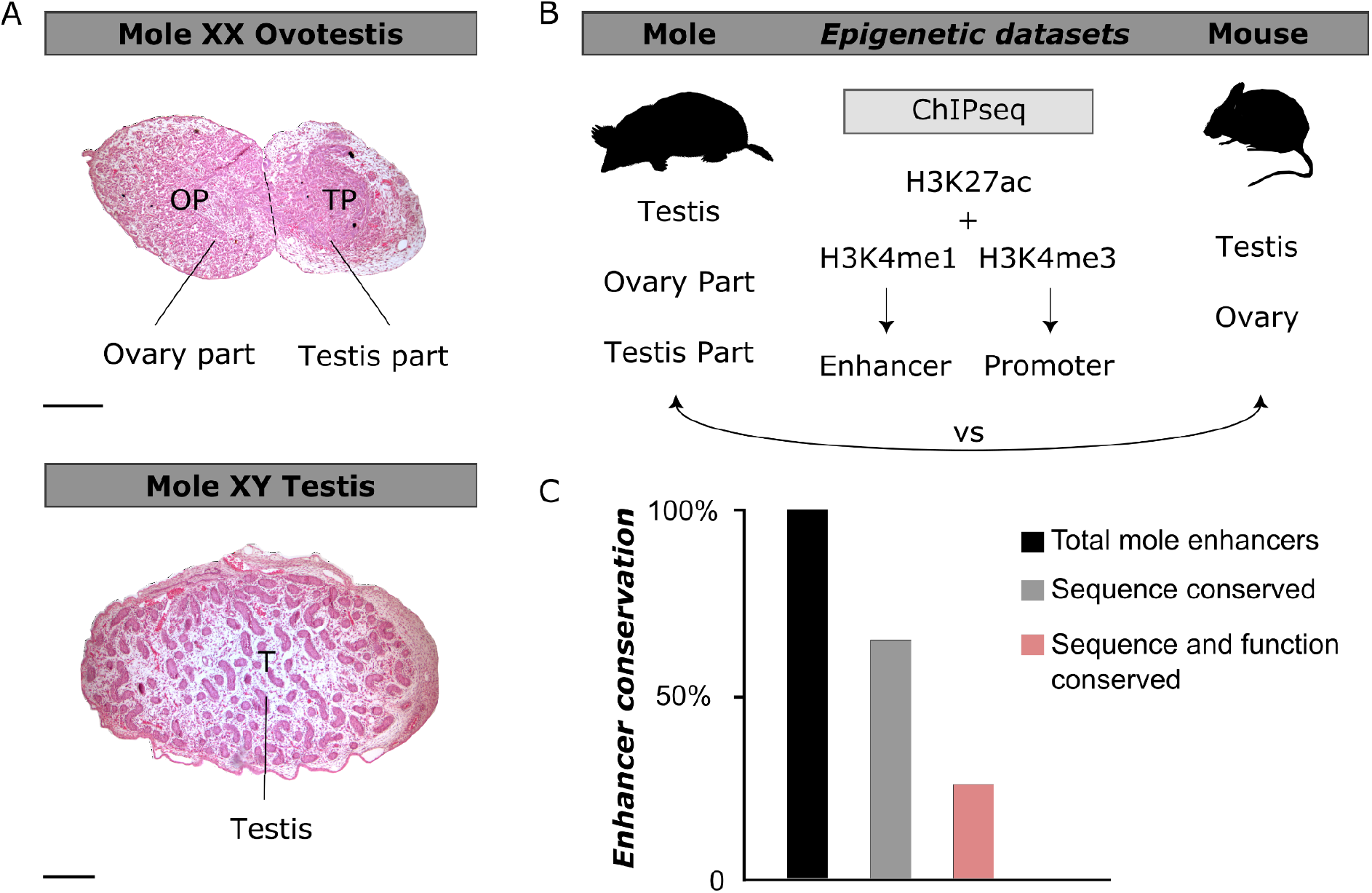
Characterization of regulatory elements in mole ovotestes. A. Hematoxylin and eosin staining of mole gonads at postnatal stage P7. Female ovotestis in upper panel, testis in lower panel. OP, TP and T means ovary part, testis part and testis respectively. Note the clear separation into two parts of the ovotestis. Scale bars represent 100μm. B. Scheme of the gonadal tissues sampled to generate the epigenetic datasets in mole and mouse. Five tissues and three different histone modifications were used for the ChIP-seq experiments. C. Percentage of mole enhancers conserved compared to mice, at the sequence level in gray and at the enhancer signature level in light red.

To explore the degree of conservation of the enhancer landscape in moles, we generated analogous datasets from mouse gonads, at a time point when Leydig cells differentiate and meiosis takes place (E13.5; **Figure 1B**). By comparing mole and mouse gonadal epigenetic datasets, we observed that from 70,618 predicted enhancers in mole gonads approximately 65% are conserved at the sequence level (**Figure 1C**). However, only 25% of those enhancers are active in both species, meaning they share an active enhancer signature in both mole and mouse gonads (**Supplementary Table 1**). Accordingly, approximately 40% of the predicted sequence conserved enhancers represent mole-specific regulatory regions and are thus potentially associated with characteristics of this species. Therefore, our results imply a prominent turnover of enhancer sequence and function during gonad evolution.

### Co-option of Sall1 expression in mole ovotestis formation

Our approach identified a subset of 6,419 mole-specific enhancers that are only active in the testicular part of the ovotestis, which could potentially contribute to the development of this unique tissue. We then explored if these enhancers are associated with the acquisition of specific transcriptional signatures using RNA-seq datasets from the same developmental stage. We therefore jointly ranked enhancers by specificity in enhancer probability in the testicular part of the ovotestes and the specific expression of their putative target gene in the same tissue. We defined the putative target genes of each enhancer as the gene with the closest transcription start site to the enhancer region within the same TAD. This approach prioritizes genes whose respective regulatory domain contains enhancer elements specifically active in the testicular part compared to the ovary part and the male testis (**Figure 2A, Supplementary Table 2**). The top-ranking genes identified by this approach were *NPY* and *SALL1. NPY* is a hormone neuropeptide expressed in Leydig cells^10,11^, whereas *SALL1* is a transcription regulator involved in cell fate decisions^12^. *SALL1* is usually expressed during development in embryonic tissues, including eye, limb or kidney^13^. Strikingly, our RNA-seq data revealed that *SALL1* is highly expressed in the testicular part of mole ovotestes at P7, but not in the XY testis or the XX ovarian region. In fact, *SALL1* is highly expressed already in the early embryonic ovotestis and becomes specific to the testis part as the organ differentiates (**Figure 2B**). In humans, mutations in *SALL1* are associated with a congenital malformation syndrome affecting limbs, kidneys and ears (Townes Brocks syndrome, OMIM #107480)^14^ and misexpression has been linked to certain types of androgen-producing ovarian tumors^15^, indicating that it might be involved in re-programming of ovarian cells.

**Figure 2.**
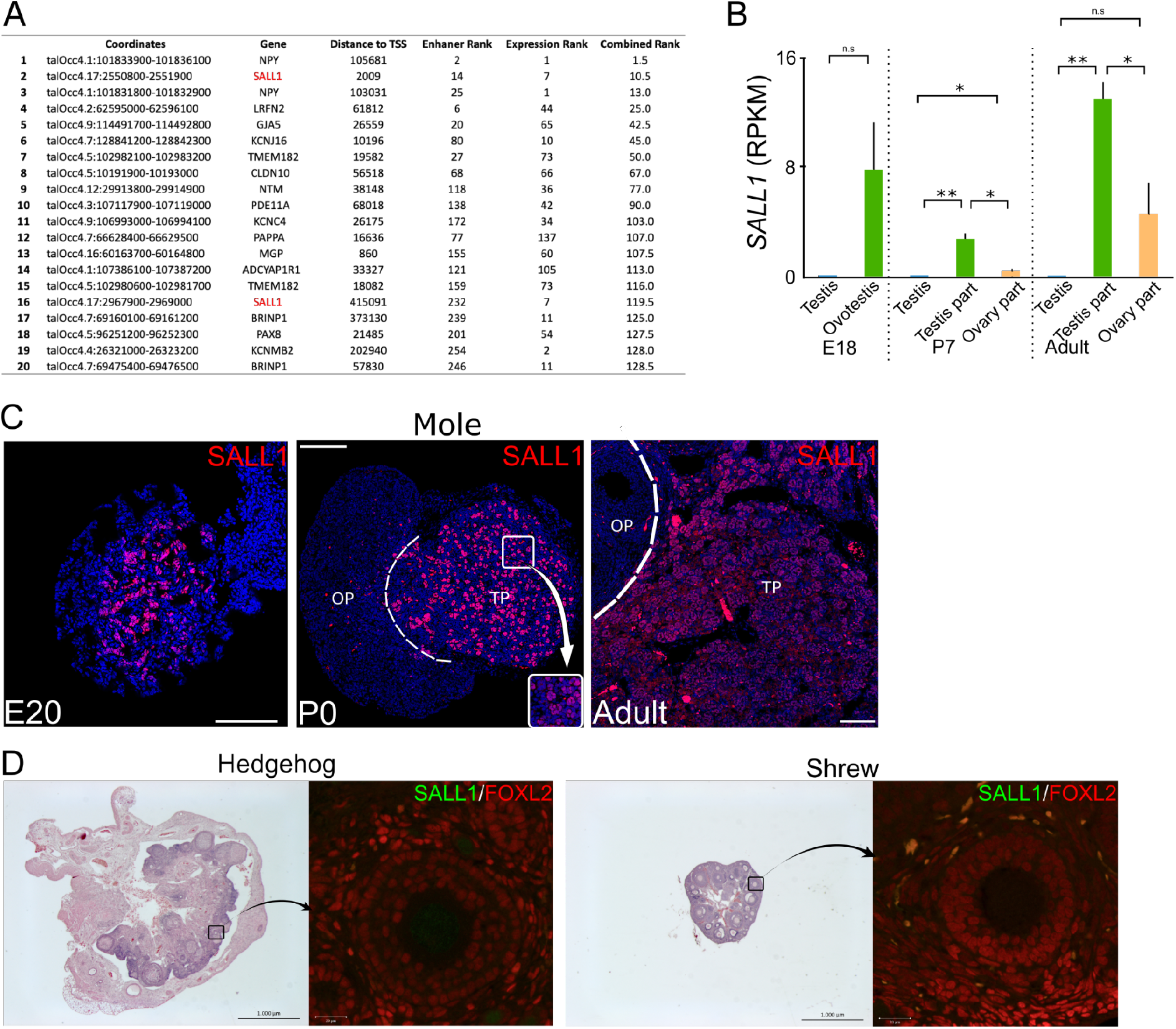
Identification of *SALL1* as a marker for testis part formation in mole ovotestes. A. Top 20 enhancer regions ranked by enhancer score and specificity of expression of the associated gene in the testis part of the ovotestis. Note the two *SALL1* enhancers which are highly ranked (rank 2+16). B. *SALL1* expression levels in RPKM from mole RNA-seq data at different developmental time points. C. Spatio-temporal profile of expression in moles (immunofluorescence, SALL1 in red; DAPI in blue). Note that SALL1 is spatially restricted to the medullary (testicular) region of the mole ovotestis at E20 and is also present in the testis part thereafter. Inset shows localization to Sertoli-like cells. OP: ovarian part, TP: testicular part. Scale bars: 100 μm. D. Spatial expression of SALL1 in adult female hedgehog (left) and adult female shrew (right) (immunofluorescence, SALL1 in green, ovarian marker FOXL2 in red). Note absence of SALL1 expression. Black and white bars represent 1000 μm and 20 μm, respectively.

To further explore the spatio-temporal dynamics of *SALL1* expression, we performed immunostaining in mole gonads at different stages of development (**Figure 2C**). This analysis revealed that SALL1 expression is specific to the mole female gonad, and importantly, this expression is spatially restricted to the medullary region of the developing ovotestis, which is the precursor of the testicular tissue. Specifically, SALL1 expression is restricted to the Sertoli-like cell population. This expression pattern is constant during the entire development and persists in adulthood, thus constituting SALL1 as a bona-fide marker for the testicular tissue of mole ovotestis.

We then explored the evolutionary conservation of SALL1 expression, by investigating the gene expression in other mammalian species. We examined the pattern of expression of *Sall1* in mice by immunostainings and transcriptomic analyses. Immunostaining analyses showed a complete absence of SALL1 protein in mouse gonads at embryonic stage E13.5, however, the protein could be detected in known *Sall1*-expressing tissues, such as the embryonic kidneys (**Supplementary Figure 1A**). This observation is extended to adulthood where RNA-seq data shows practically no expression in both males and females when compared to the mole (**Supplementary Figure 1B**). We further expanded our analysis of SALL1 expression to also include species from the order *Eulipotyphla*, which are evolutionarily close to moles^16^. Specifically, we analyzed ovarian samples from the hedgehog *Atelerix albiventris*, as well as from the common shrew, *Sorex araneus*, the latter species belonging to the closest taxonomic group but developing normal ovaries. Immunostaining analyses showed an absence of SALL1 expression in the gonads of these two species (**Figure 2D**). Overall, these results suggest that *SALL1* expression has been co-opted in mole ovotestis development.

### Conserved 3D organization but divergent enhancers at the mole SALL1 locus

To define the regulatory landscape of *SALL1*, we examined previously published Hi-C data from different mole tissues^7^ (**Figure 3A, Supplementary Figure 2**). Chromatin interaction maps revealed a large 1 Mb TAD, in which *SALL1* is the only protein-coding gene. The interaction profile of *SALL1* in the testicular part of the ovotestis was further explored at increased resolution through circular chromosome conformation capture (4C-seq), using the gene promoter as a viewpoint (**Figure 3B**). These experiments demonstrate prominent interactions of *SALL1* across the entire TAD, with a sharp decrease in contacts outside this domain. We then explored the degree of conservation of the *SALL1* interaction profile by comparing the mole against mouse data^17^. This comparison revealed that, despite notable differences in *SALL1* expression, the locus displays a remarkable preservation of the 3D structure across species (**Supplementary Figure 3A**).

**Figure 3.**
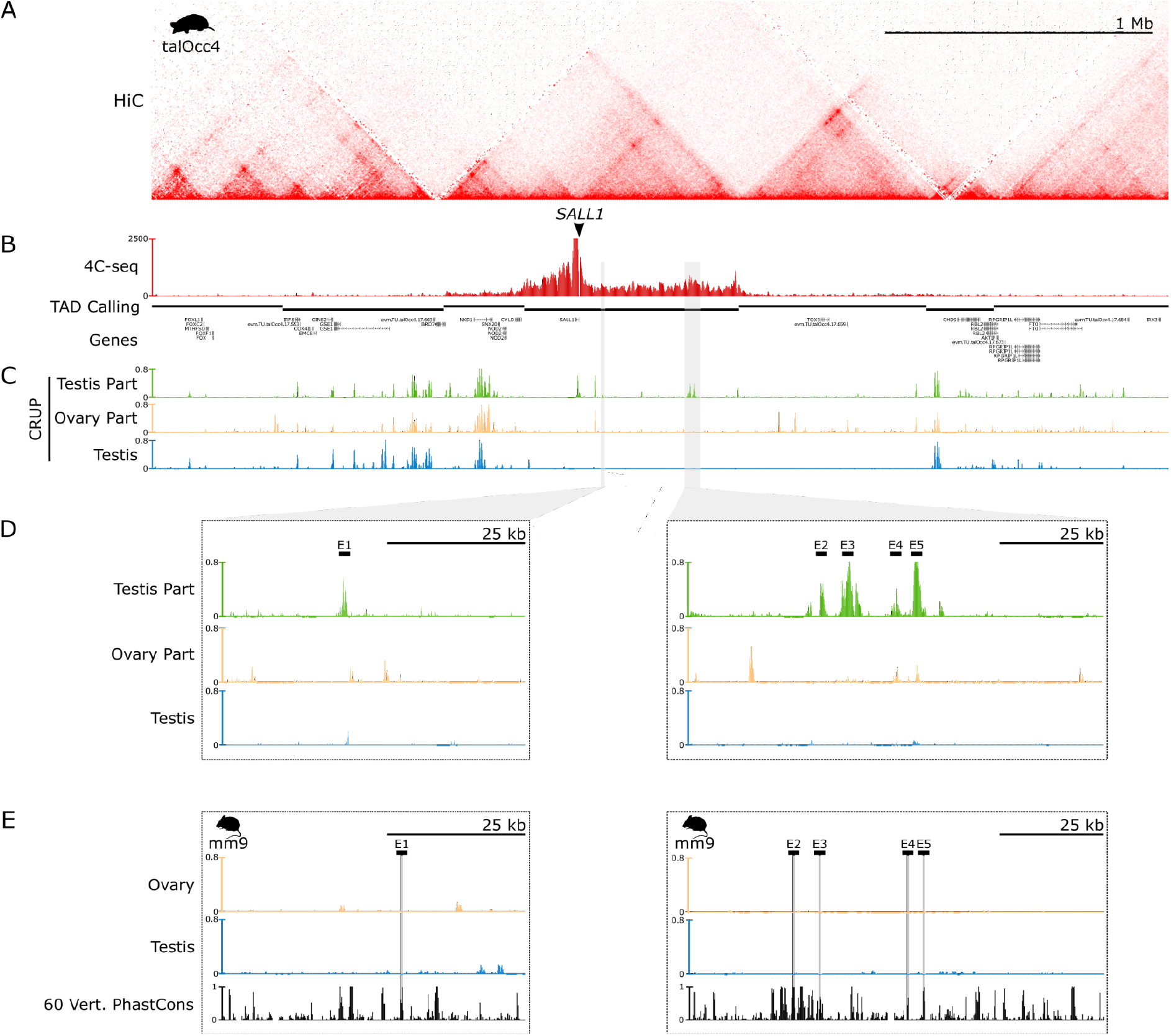
Regulatory domain and the epigenetic landscape of *SALL1*. A. HiC map from mole embryonic limbs denotes the domain of *SALL1* in a large gene desert. B. 4C-seq analysis from female adult testis part with *SALL1* as viewpoint. Note high interaction frequency between the gene promoter and the surrounding 1Mb desert clearly demarcating the *SALL1* regulatory domain. C. Epigenetic landscape of *SALL1* in the three tissues sampled, called with the tool CRUP. Note the presence of numerous active enhancers in the testicular part of the ovotestis where *SALL1* is specifically expressed. D. Zoom in on two mole regions containing five specific regulatory elements for the testis part of the ovotestes, named as E1-5. E. Homologous regions to the testis part enhancers in the mouse genome. Homologous regions are marked as gray bars. Note the absence of enhancer activity of these regions despite the sequence conservation in vertebrates ^18^(60 vertebrates Basewise Conservation by PhyloP).

Next, we overlaid the *SALL1* interaction profile with the epigenetic datasets, to identify potential regulatory elements (**Figure 3C**). This revealed several candidate enhancer regions that were active in the testicular part of the ovotestis. Specifically, we identified one enhancer unique for the testicular portion close to *SALL1*, as well as a distant cluster of four enhancers. This enhancer cluster is indeed in close physical proximity to the *SALL1* promoter, as denoted by a specific loop in the Hi-C map and an increase in contacts in the 4C profile. A zoom in on these regions highlights the specificity of these enhancers for the testicular part of the ovotestes (**Figure 3D**). A comparison with the respective mouse epigenetic datasets revealed that these elements were not active in mouse gonads but nevertheless show a high degree of sequence conservation across vertebrates. This conservation at the sequence level denotes the potential of these regions to evolve at the regulatory level (**Figure 3E**).

To validate the activity of these putative enhancers *in vivo*, we tested the mole regions for enhancer activity in mouse transgenic LACZ reporter assays^19^. We selected the five mole elements (E1-5; **Figure 3D**) that display high conservation across vertebrates but are not functionally conserved in mouse gonads. At E13.5, all of the tested regions showed reproducible tissue-restricted activity, thus confirming them as true enhancers (**Figure 4; Supplementary Figures 4-8**). Enhancer activity was observed in several tissues, such as the limbs or eyes, in which *Sall1* is known to be expressed. Interestingly, enhancer 3 displayed specific activity in kidneys, another *Sall1*-expressing tissue^13^, which is consistent with its predicted enhancer activity in this tissue (**Supplementary Figure 3B**). While none of these enhancers induced reporter expression in developing gonads, enhancers 1, 2, 4 and 5 were active in the adjacent mesonephros. Importantly, this tissue has the same ontogenetic origin as the gonads, and contributes to its cellular composition, through cell migration. These enhancers are possibly activated in mole ovotestes by a different pool of transcription factors, not present in mouse gonads. However, their mesonephric activity in mice and their vertebrate conservation suggests a prominent evolvability for these elements. Altogether, our results suggest that the evolution of enhancers may underlie *SALL1* expression in the testicular part of the mole ovotestis.

**Figure 4.**
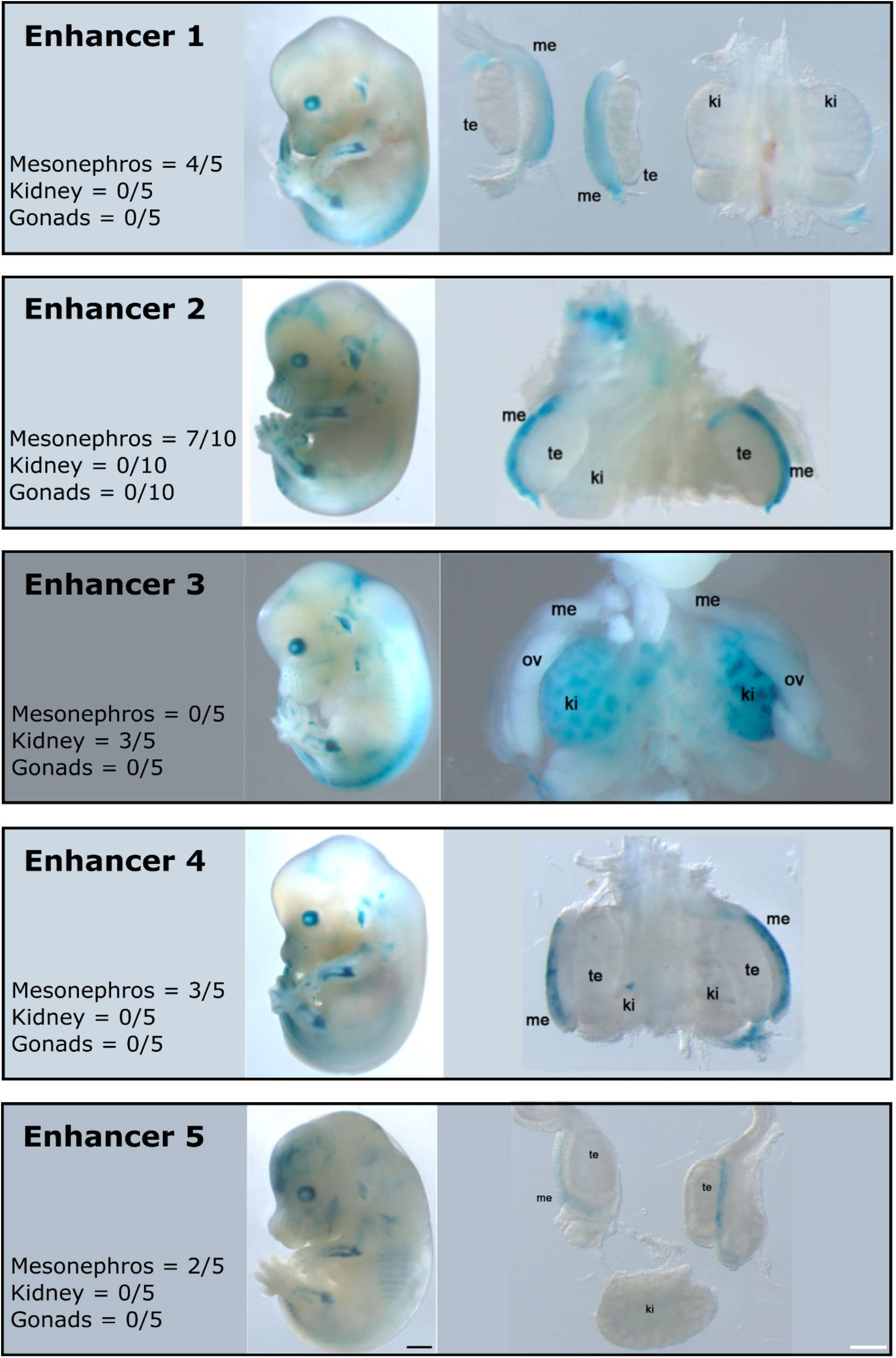
LACZ reporter assays for enhancer elements E1-5 associated with *SALL1*. The enhancer activity of each element is depicted in a separated box 1 to 5. Entire embryos at E13.5 are displayed next to the dissected urogenital tracts. Me: mesonephros, te: testes, ov: ovaries, ki: kidneys. Black and white bars represent 1000 and 100 μm, respectively.

### SALL1 expression triggers specific gene expression programs during ovarian development

To investigate the effects of *Sall1* expression during early gonadal development, we induced its expression in the mouse ovary. For this purpose, we created a BAC construct to overexpress *Sall1* in somatic ovarian cells (**Figure 5A**). The BAC contains the regulatory elements and the promoter of the *Wt1* gene, which is constitutively expressed in gonadal somatic cells^20^, but the gene is replaced by the coding sequence of *Sall1*. Through *PiggyBac* transgenesis, we integrated this construct into female mouse embryonic stem cells (mESC), which were subsequently used to generate transgenic mice through morula aggregation. In contrast with the wildtype ovaries, mutant ovaries successfully express *Sall1* in the somatic cells, as denoted by the overlapping signal with *Foxl2*, a bonafide marker for female somatic ovarian cells^21^ (**Figure 5B**). However, at the phenotypic level, adult female mice did not show major morphological gonadal alterations and breed normally. This suggests that *Sall1* alone is not sufficient to induce the development of testicular structures.

**Figure 5.**
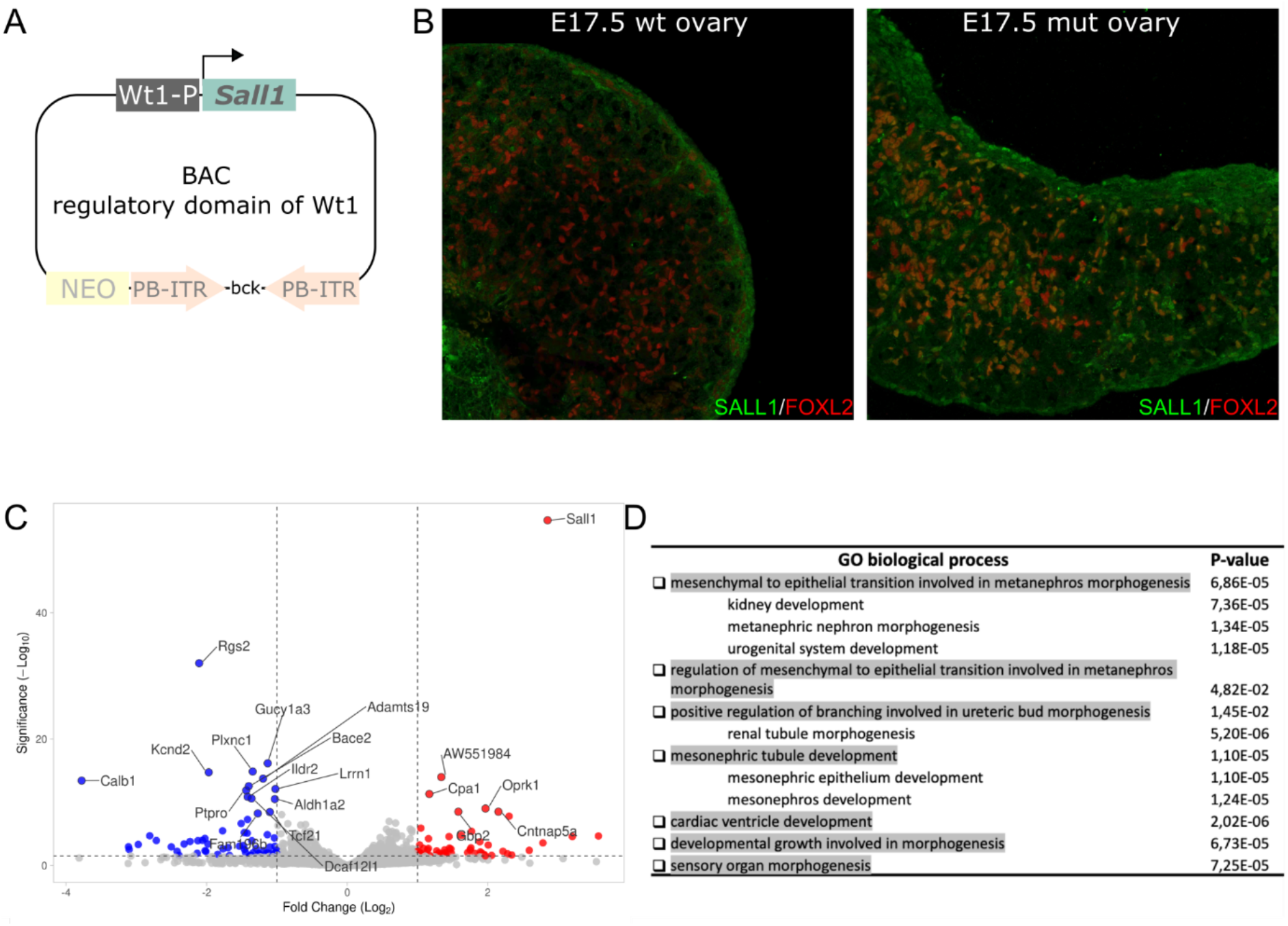
Overexpression of *Sall1* in mouse embryonic ovaries results in hundreds of dysregulated genes. A. Cloning strategy to overexpress *Sall1* in somatic ovarian cells through BAC transgenesis. *Sall1* is regulated under the promoter and regulatory regions of the gonadal somatic gene, *Wt1*. B. Immunostainings against SALL1 (green) and FOXL2 (red) in wildtype and mutant ovaries at E17.5. Note the high abundance of double positive cells (orange) for SALL1 and FOXL2 in the mutant gonad, confirming the overexpression success. C. Volcano plot from RNA-seq of mutant ovaries compared to control ovaries from littermates at E13.5. Names of the 20 most deregulated genes are highlighted. Note that *Sall1* is the most significantly upregulated gene. D. Gene ontology enrichment analyses of the common upregulated genes in the *Sall1* mutant ovaries and in the testis part of the ovotestes.

To gain further insights into the molecular signatures of *Sall1* ovarian expression, we performed RNA-seq in gonads from mutants and litter-mate controls at E13.5. This analysis revealed around 400 deregulated genes where *Sall1* is the most significantly up-regulated gene (**Figure 5C, Supplementary Table 3**). To understand the consequences of *Sall1* expression in female gonads, we compared the deregulated genes in the mutant ovaries to those specifically expressed in the testis part of the ovotestis. We found 56 and 36 genes commonly upregulated and downregulated, respectively, in the mutant mice and in the female testicular part in moles. Gene ontology enrichment analyses reveal no enrichment for the downregulated genes, however the upregulated genes were enriched in terms related to kidney development, ureteric bud morphogenesis and mesonephros development (**Figure 5D, Supplementary Figure 9**). The upregulation of genes involved in mesonephros development is concordant with the reporter activity of the mole testis part enhancers in the mesonephros of mice. These results showed that similar gene regulatory networks are active in the mesonephros and in the testis part of the ovotestis and highlight that the SALL1 gene co-opted for ovotestes development through the acquisition of specific enhancers.

## DISCUSSION

Across vertebrates, gonadal development is characterized by a remarkable evolutionary plasticity^22,23^. This is particularly highlighted by the development of ovotestes in moles, in which the development of a testicular region that increases the production of male hormones is fully compatible with a reproductive function^6^. In previous studies, we demonstrated that mole ovotestis development is associated with a prolonged expression of *FGF9* through early gonadal development^7^. This heterochronic expression pattern delays the onset of female meiosis and creates a pro-testicular environment that is critical for ovotestis development. Our transgenic experiment revealed that *SALL1* overexpression can activate ectopic gene expression programs. Interestingly, this program is characterized by molecular signatures that are shared with other *SALL1*-expressing tissues, including kidneys, and that affect the expression of mesonephros-related genes. This program is not sufficient to trigger sex-reversal mechanisms, as denoted in phenotypical analyses. Therefore, it is plausible that *SALL1* may cooperate with other factors in ovotestis development and/or benefit from the pro-testicular environment that *FGF9* misexpression induces.

During evolution, genes are frequently co-opted for species-specific processes. These effects are often mediated by changes in the activity of regulatory elements that preserve the essential function of genes and, at the same time, allow a diversification of its expression in new tissues and cell types^24–26^. Mole enhancers were not able to recapitulate gonadal expression in mouse reporter assays. This might indicate that additional trans-acting factors are required for their activation. However, *SALL1* enhancers also display consistent activity in the mesonephros, a tissue that shares a common molecular origin with the gonad. Furthermore, the mesonephros is a known source of endothelial, myoid and supporting cells to the gonad^27,28^. Interestingly, the developing ovotestes of the mole, in contrast to female gonads of most mammalian species, show a prominent expression of migration markers^29,30^. Thus, migrating cells from the mesonephros could contribute to the mole ovotestis. In fact, the overexpression of Sall1 in ovaries activates genes enriched in mesonephros development and these genes are shared with the testis part of the ovotestis. Thus, our finding suggests that the origin of this tissue might be the mesonephros with *SALL1* being co-opted to initiate common pathways. The mesonephric activation of SALL1 is driven by several enhancers, resembling a functional redundancy of CREs that has been described at multiple developmental loci^31^. Such cooperative activity has been proposed to arise by an initial gain of transcription factor binding sites that is progressively stabilized through the recruitment of additional sites at other elements, giving capacity to these elements to evolve in different functional directions^32^.

TAD structures serve as a spatial scaffold, in which regulatory elements interact with their cognate genes, thus representing the existence of large 3D regulatory landscapes contributing to the specificity of gene expression. These domains have been suggested to represent a fertile ground for the evolution of gene expression^33–35^. Previous studies have demonstrated that TADs impose important constraints during evolution, as genomic rearrangements are more prone to occur at boundaries, preserving TADs as entire regulatory units^36^. However, genomic rearrangements that reorganize TADs can be also associated with changes in gene expression that might induce the evolution of traits^37^. This has been recently exemplified with the ectopic activation of the PCP pathway, linked to the development of enlarged fins in skates but also in moles where genomic rearrangements affecting the *FGF9* and *CYP17A1* TADs are associate to intersexuality^7,38^. The results of these studies also highlight that the evolution of CREs within conserved TADs is a prominent mechanism for evolution. This is exemplified by the striking conservation of TAD organization at the *SALL1* TAD, which is characterized by a remarkable internal evolution of CREs. These results are consistent with previous observations and further reinforce the idea that TADs might serve as a scaffold for the evolution of gene pleiotropy^26,39^. In summary, our results demonstrate the co-option of *SALL1* in mole ovotestis development, through regulatory changes that occur in spite of a striking conservation of TAD organization. This highlights the multilayered nature of gene regulation and how changes at different levels can serve as a driving force for the evolution of traits.

## Supplementary figures

**S1.**
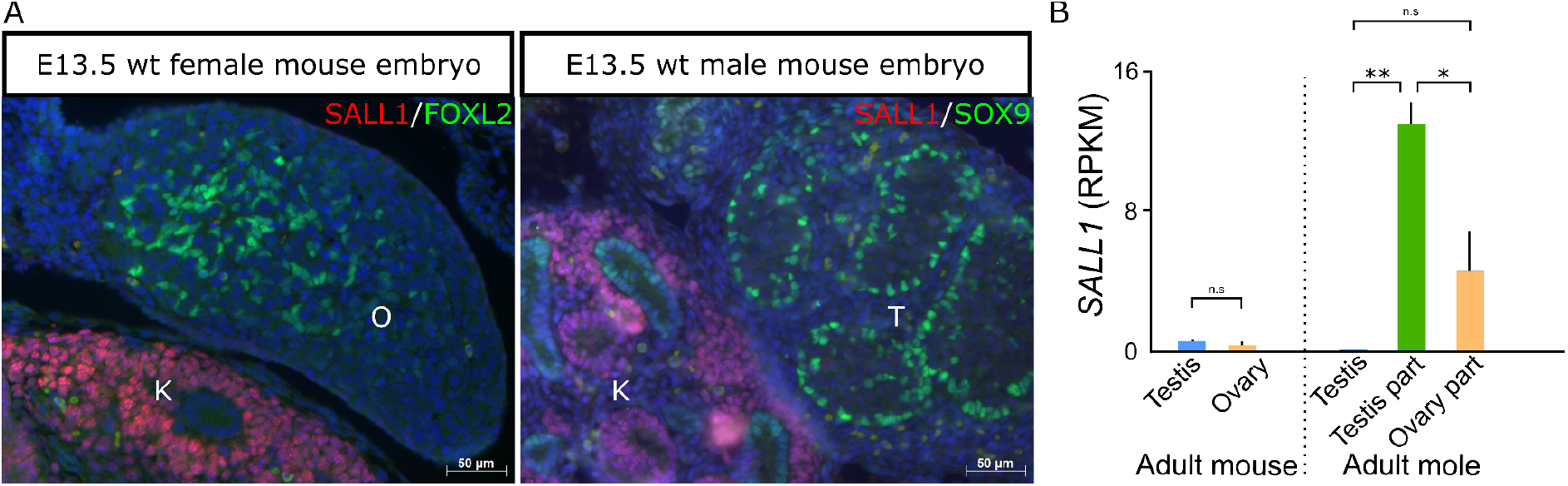
*Sall1* expression pattern in mouse gonads. A. Immunostainings of SALL1 (red) and FOXL2 and SOX9 (green) as markers of somatic female and male cells, respectively. O: ovary, T: testis, K: kidney. Note the absence of SALL1 positive cells in the embryonic gonads but the specific expression in the adjacent kidney. B. RPKMs quantification from RNA-seq data of adult gonads in mouse and mole. Expression levels in mouse are lower compared to mole and not sex-specific.

**S2.**
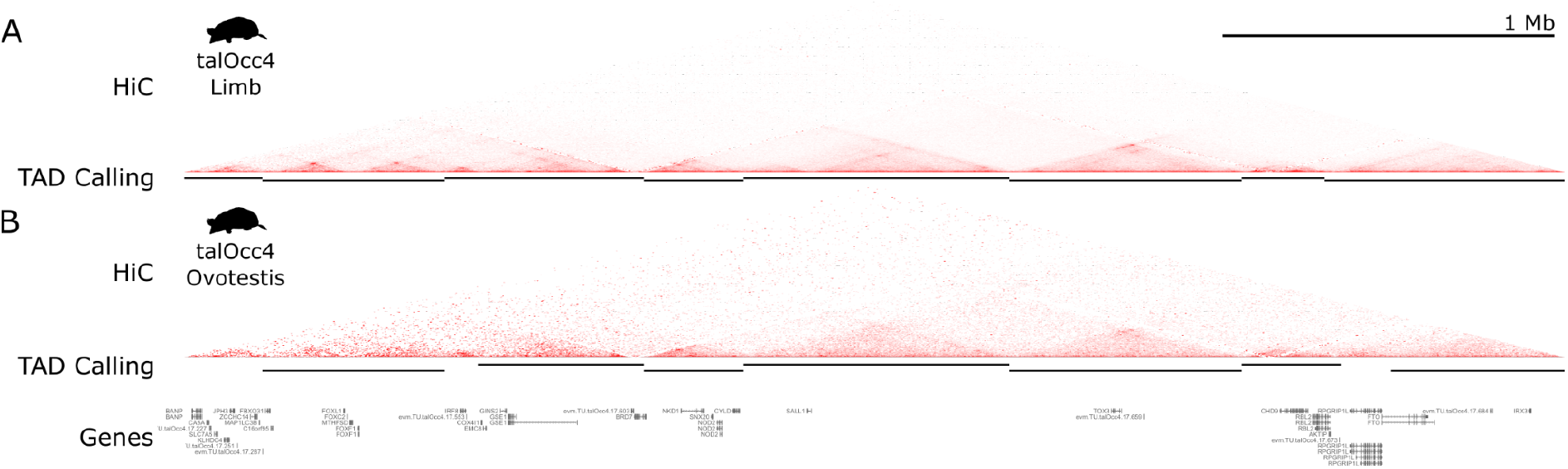
HiC maps comparison between limb and ovotestis. A. HiC maps at high resolution from embryonic limbs with the corresponding TAD calling (black bars) underneath. B. HiC maps from adult ovotestis with the corresponding TAD calling (black bars) underneath. Note the conservation of the *SALL1* TAD domain between tissues.

**S3.**
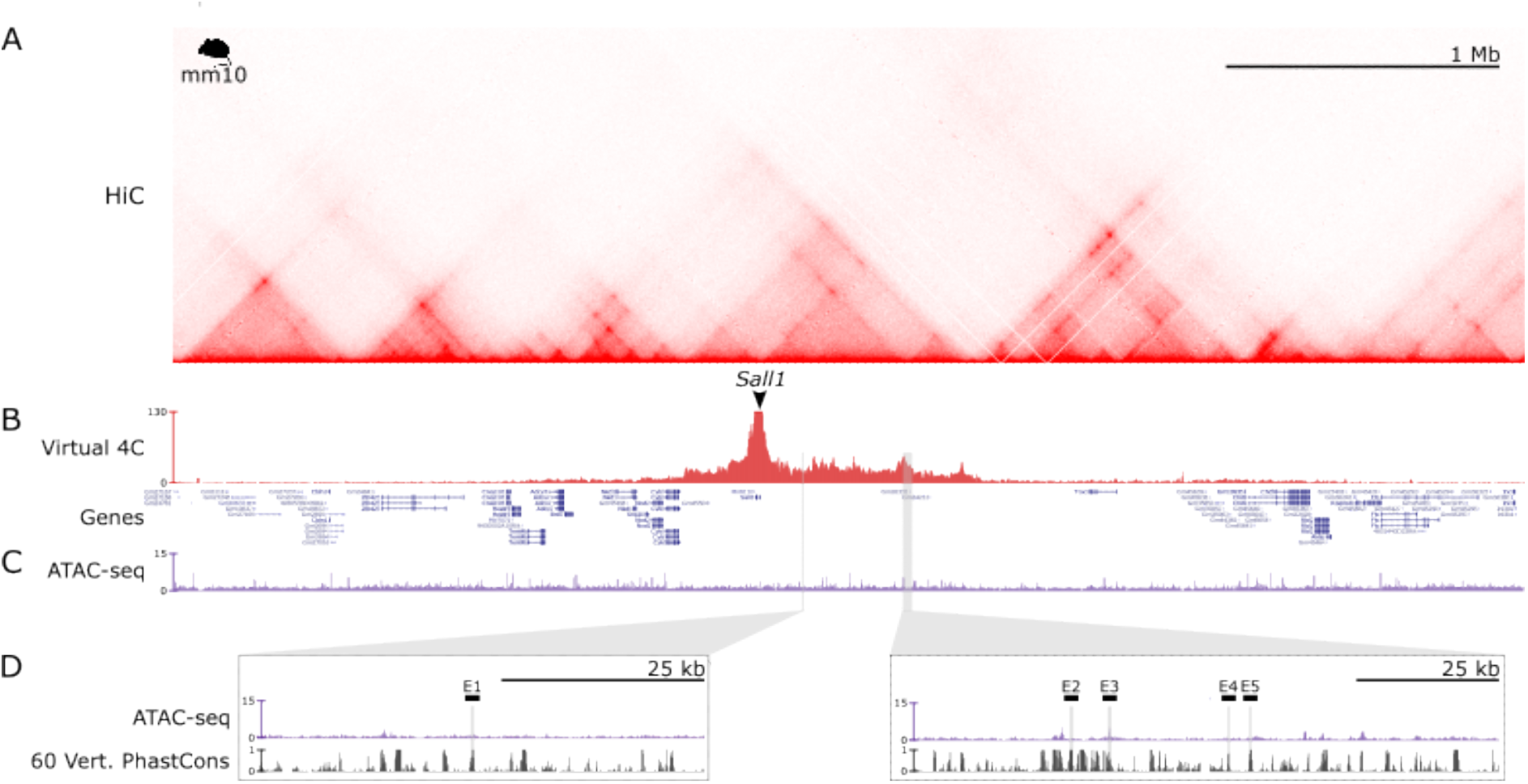
Regulatory domain of *Sall1* in mouse. A. HiC map from Neural progenitor Cells (NPCs) denotes the domain of *Sall1* in a large gene desert. B. Virtual 4C-seq analysis from NPCs HiC maps with *SALL1* as viewpoint. Note high interaction frequency between the gene promoter and the surrounding 1Mb desert clearly demarcating the *Sall1* regulatory domain. The domain is strikingly conserved between cell types and species. C. ATAC-seq track from mouse embryonic kidneys at E14.5 to identify regulatory regions in this tissue. D. Zoom in on the two equivalent regions where the mole enhancers were identified. Homologous regions are marked as gray bars and labeled as E1-5. Consistent with our enhancer activity results, enhancer 3 (E3) coincides with an ATAC-seq peak in kidneys.

**S4.**
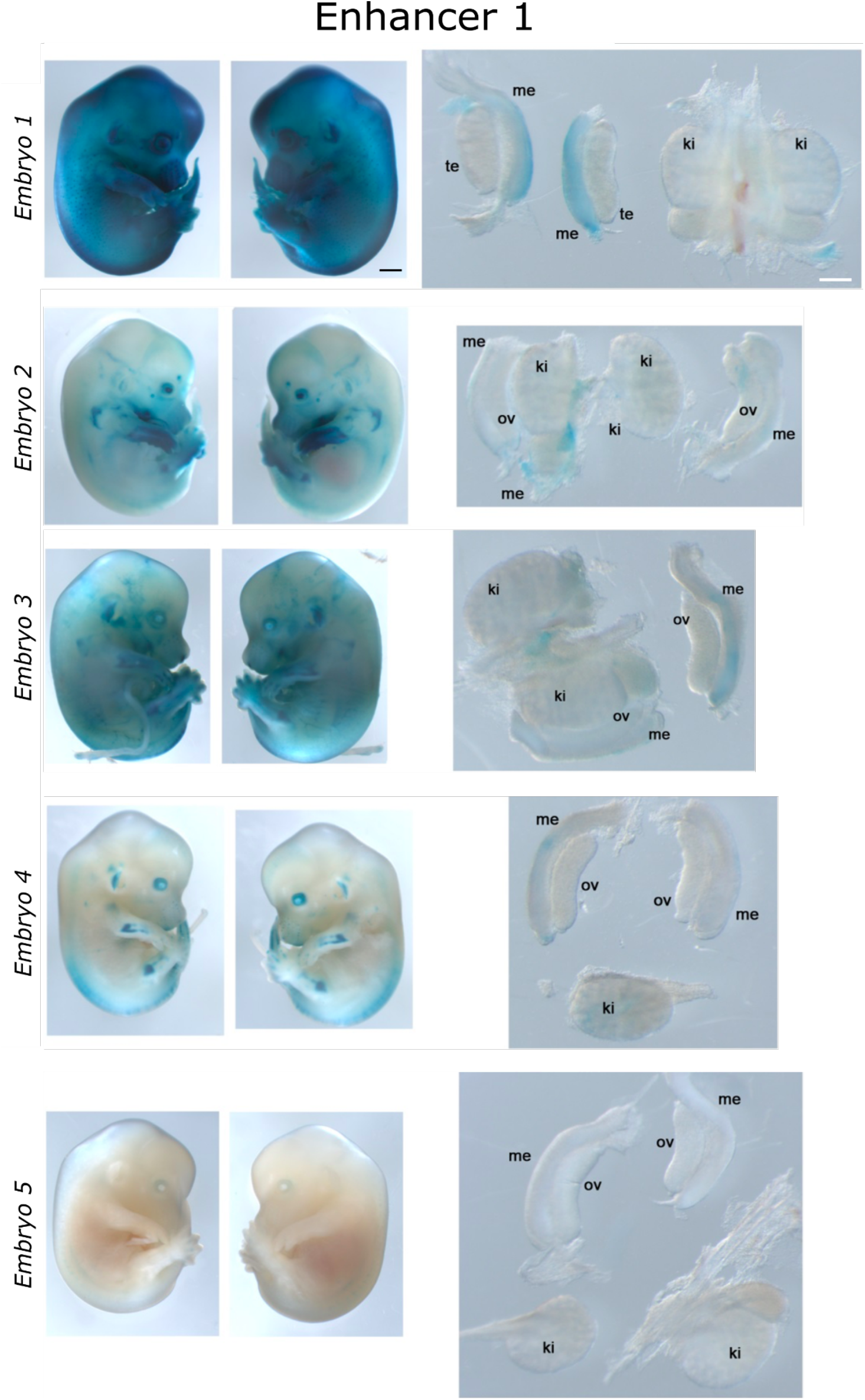
LACZ enhancer reporter assay for Enhancer 1. All embryos analyzed for this enhancer are depicted. Entire embryos at E13.5 are displayed next to the dissected urogenital tracts. Me: mesonephros, te: testes, ov: ovaries, ki: kidneys. Four out of five embryos showed mesonephros-specific staining. Black bars: 1000 μm, white bars: 100 μm.

**S5.**
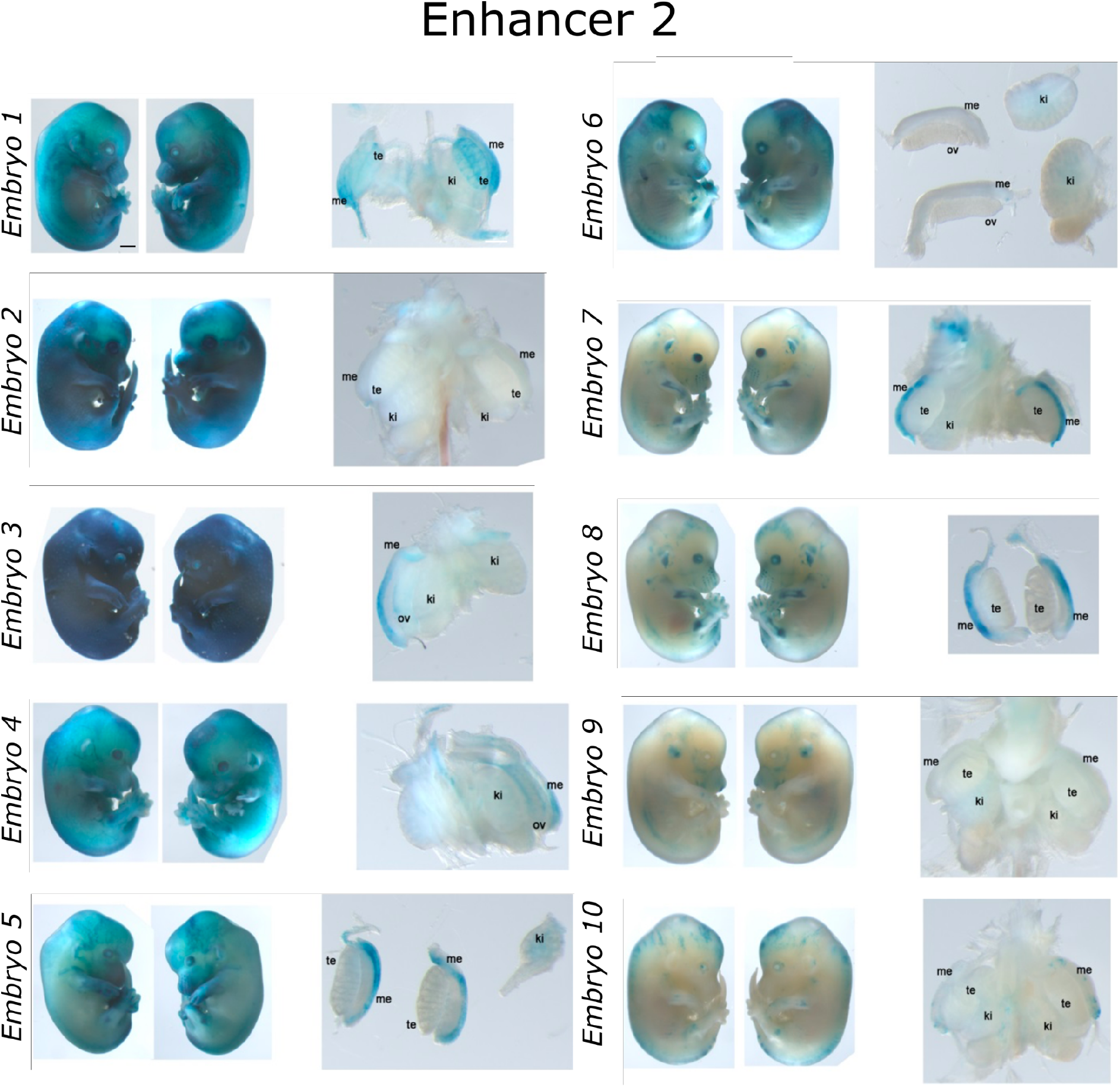
LACZ enhancer reporter assay for Enhancer 2. All embryos analyzed for this enhancer are depicted. Entire embryos at E13.5 are displayed next to the dissected urogenital tracts. Me: mesonephros, te: testes, ov: ovaries, ki: kidneys. Seven out of ten embryos showed mesonephros-specific staining. Black bars: 1000 μm, white bars: 100 μm.

**S6.**
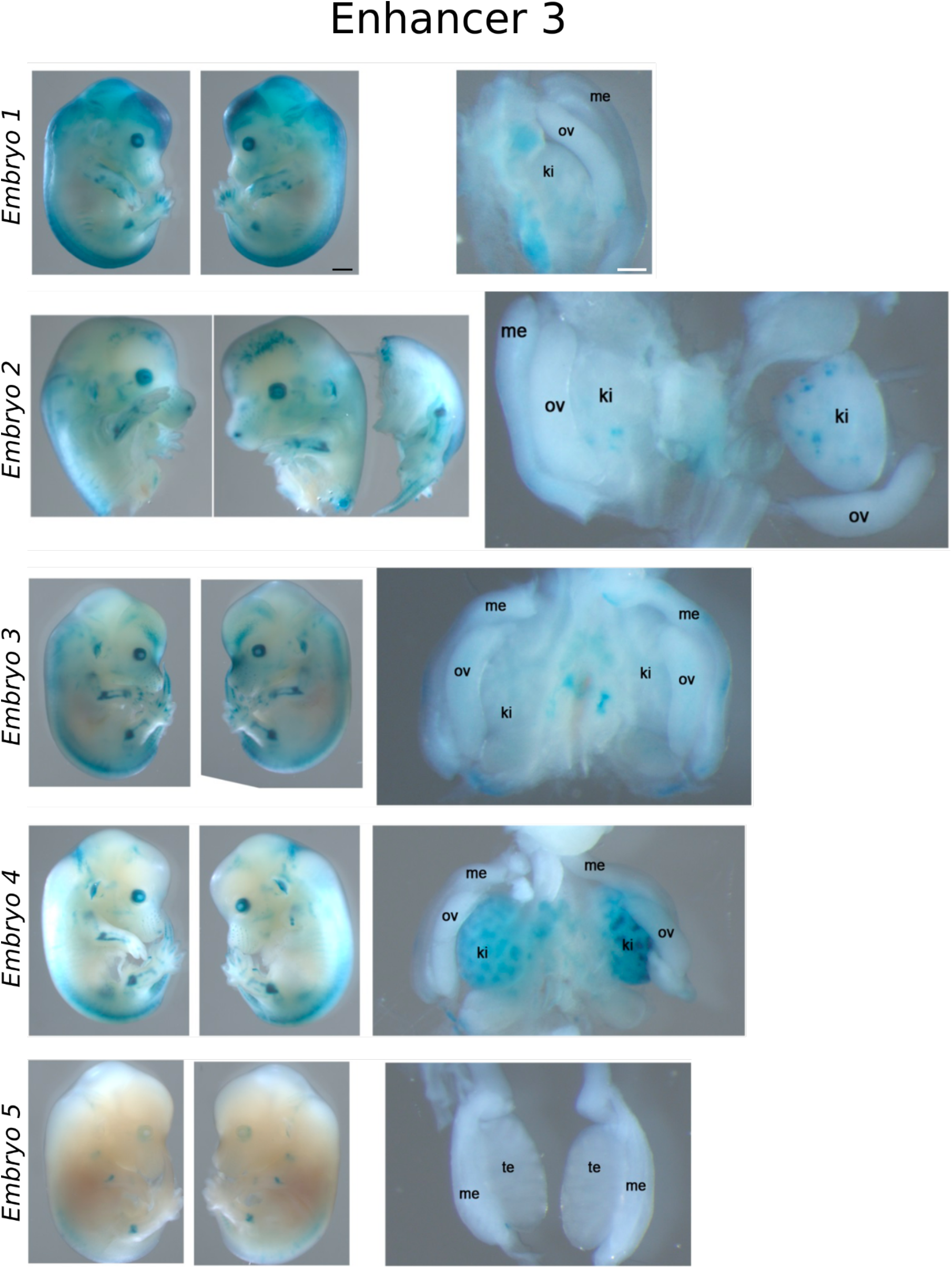
LACZ enhancer reporter assay for Enhancer 3. All embryos analyzed for this enhancer are depicted. Entire embryos at E13.5 are displayed next to the dissected urogenital tracts. Me: mesonephros, te: testes, ov: ovaries, ki: kidneys. Three out of five embryos showed kidney-specific staining. Black bars: 1000 μm, white bars: 100 μm.

**S7.**
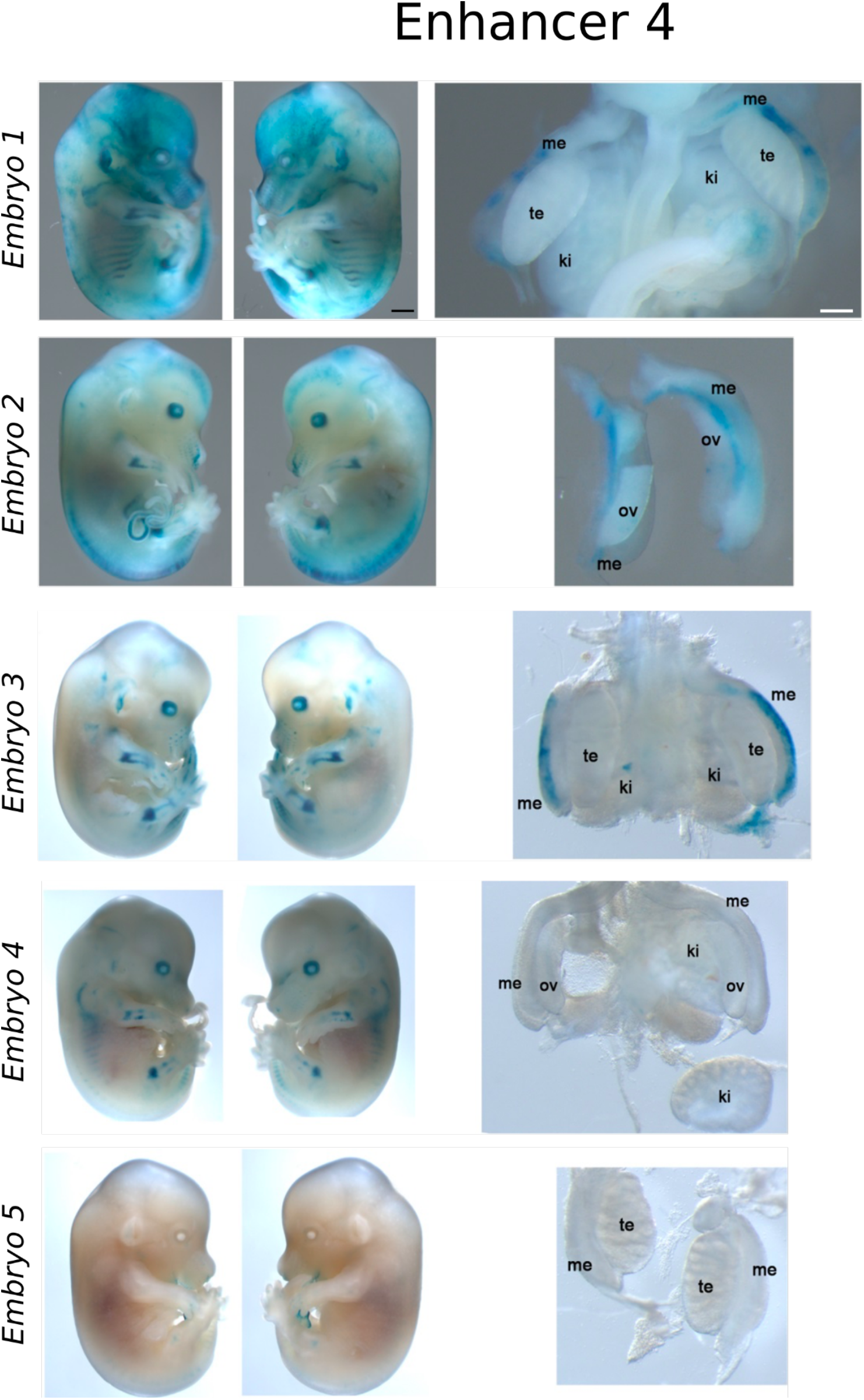
LACZ enhancer reporter assay for Enhancer 4. All embryos analyzed for this enhancer are depicted. Entire embryos at E13.5 are displayed next to the dissected urogenital tracts. Me: mesonephros, te: testes, ov: ovaries, ki: kidneys. Three out of five embryos showed mesonephros-specific staining. Black bars: 1000 μm, white bars: 100 μm.

**S8.**
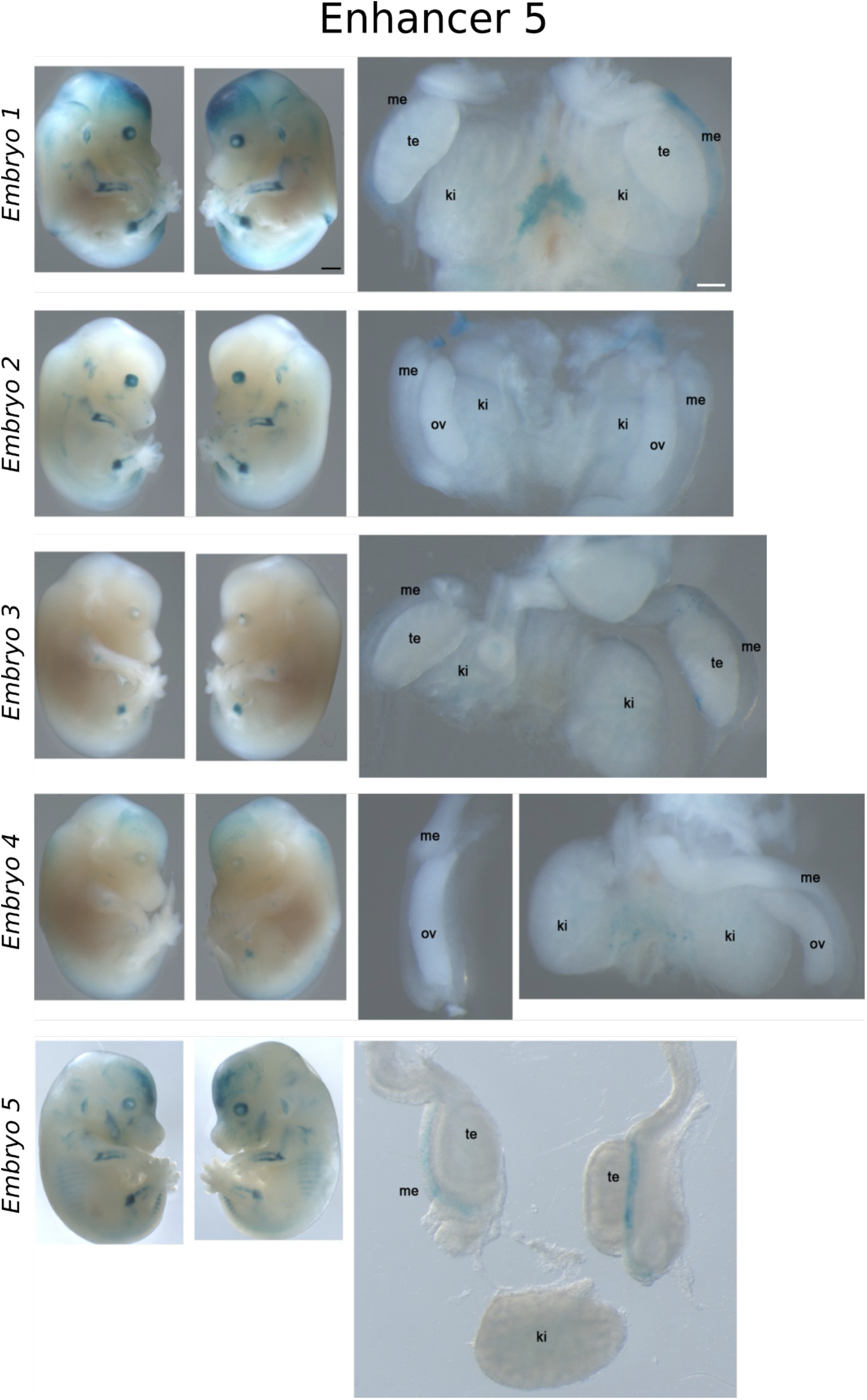
LACZ enhancer reporter assay for Enhancer 5. All embryos analyzed for this enhancer are depicted. Entire embryos at E13.5 are displayed next to the dissected urogenital tracts. Me: mesonephros, te: testes, ov: ovaries, ki: kidneys. Two out of five embryos showed mesonephros-specific staining. Black bars: 1000 μm, white bars: 100 μm.

**S9.**
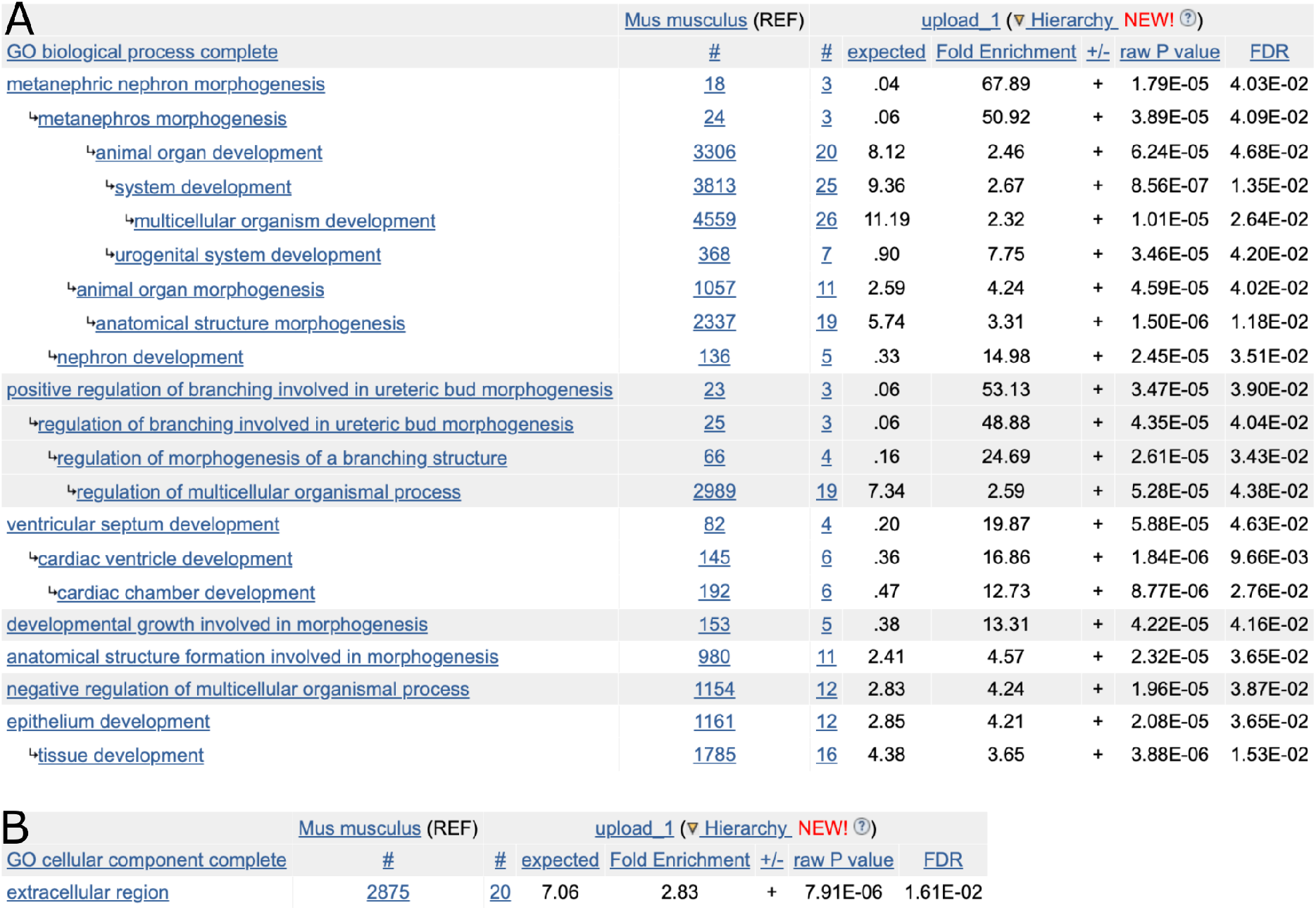
Gene ontology enrichment of commonly upregulated genes in female mole testis part and mouse *Sall1*-overexpressing mutants. A. GO terms for biological processes. B. GO terms for cellular components.

## MATERIAL AND METHODS

### Animal models

Adult, infant or embryonic specimens of the Iberian mole *Talpa occidentalis* were used with annual permission from the Andalusian Environmental Council. The animals were captured alive in poplar groves plantations in Santa Fe, Chauchina and Fuentevaqueros (Granada province, southern Spain) using an efficient trapping system as described in a previous publication^40^ and handled according to the guidelines and approval of the “Ethical Committee for Animal Experimentation” of the University of Granada.

Hedgehogs (*Atelerix albiventris*) were maintained in the LANE animal facility at the University of Geneva and were sampled under the experimentation permit GE24/33145 approved by the Geneva cantonal veterinary authorities, Switzerland.

Shrews (*Sorex araneus*) were trapped in wooden traps and euthanized with an isoflurane overdose followed by open-heart perfusion (see^41^ for details) in Möggingen, Germany, under permit number 35-9185.81/G-11/21 to Dina Dechmann.

LacZ transgenic mice were carried out at Lawrence Berkeley National Laboratory (LBNL, CA, USA) was reviewed and approved by the LBNL Animal Welfare Committee. Transgenic mice were housed at the Animal Care Facility (the ACF) at LBNL. All transgenic experiments were performed in accordance with national laws and approved by the national and local regulatory authorities. All animal work at Mice were monitored daily for food and water intake, and animals were inspected weekly by the Chair of the Animal Welfare and Research Committee and the head of the animal facility in consultation with the veterinary staff. The LBNL ACF is accredited by the American Association for the Accreditation of Laboratory Animal Care International (AAALAC).

The experiments for Sall1 overexpression transgenic mice were performed as approved by LAGeSo Berlin under license numbers G0346/13 and G0247/13. Transgenic experiments were performed using mouse embryonic stem cells (mESCs) from a C57BL/6J or C57BL/6J-129 hybrid background. For RNA-seq and ChIP-seq experiments, gonads from wild-type CD1 mice were used.

### Histological and immunostaining analyses

Gonads from adult animals, infants and embryos were fixed in 4% PFA and embedded in paraffin. The embedded samples were sectioned in 5μm thick slides and stained with hematoxylin-eosin according to standard protocols.

For protein spatio-temporal detection experiments, indirect immunofluorescence was used. In brief, sample slides were incubated overnight with the primary antibody at a dilution according to the manufacturer’s instructions. Next, samples were incubated with specific Alexa secondary antibodies 488 and 568 together with DAPI for one hour at room temperature. Slides were then mounted in fluoromount-G solution (SouthernBiotech) and pictures were taken either with a Laser confocal Zeiss LSM700 or Zeiss Axiovert 200M microscope.

Primary antibodies and working dilutions are listed here: mouse anti-SALL1 (abcam ab41974, dilution 1:100), goat anti-FOXL2 (abcam ab5096, dilution 1:200), rabbit anti-FGF9 (Santacruz sc-7876, dilution 1:75), rabbit anti-SCP3 (abcam ab15093, dilution 1:200), rabbit anti-SOX9 (Cell Signaling #82630, dilution 1:200), rabbit anti-CYP17A1 (kindly provided by Prof. A. J. Conley^42^, dilution 1:400) and mouse anti-CYP19A1 (abD Serotec, MCA2077S dilution 1:50).

### ChIP sequencing

Gonads from P7 moles and from E13.5 mouse embryos were fixed using 1% formaldehyde and subsequently snap-frozen and stored at −80°C. Chromatin immunoprecipitations were performed using the iDeal ChIP-seq Kit for Histones (Diagenode, Cat. No. C01010051) according to the manufacturer’s instructions. Briefly, whole fixed gonads were lysed and subsequently sonicated using a Bioruptor (45 cycles, 30 seconds on, 30 seconds off, at high power) in the provided buffers. 5 μg of sheared chromatin were then used per immunoprecipitation with 1 μg of the following specific histone antibodies: anti-H3K4me3 (Millipore, cat. No. 07-473), anti-H3K4me1 (Diagenode, cat. No. C15410037), anti-H3K27ac (Diagenode, cat. No. C15410174), and anti-H3K27me3 (Millipore, cat. No. 07-449). The samples were sequenced using Illumina HiSeq technology according to standard procedures. Mapping was performed with the STAR v2.6.1d software^41^ using settings to enforce unspliced read mapping (--alignEndsType EndToEnd -- alignIntronMax 1 --outFilterMatchNminOverLread 0.94). Finally, deduplication was performed via bamUtil (version 1.0.14; option –rmDups, https://github.com/statgen/bamUtil/releases)

### Enhancer calling and conservation

Calling of putative enhancer regions was performed for mole and mouse via the software CRUP with replicates merged beforehand. Enhancer regions with a distance <=200bp were merged. To reduce outlier effects in enhancer probability scores, a smoothing over 5 bins of 100bp was applied. In line with the original CRUP results, the probability of an enhancer region is defined as the, now smoothened, maximum score of the 100bp bins overlapping the enhancer. For the analysis of enhancer conservation, mole enhancer regions were lifted-over to the mouse genome (mm9). By definition, only those regions overlapping a conserved sequence block can be lifted, therefore depending on genome alignment settings. Here, we performed a sensitive pair-wise one-to-one genome alignment using LAST with automated training of optimal alignment parameters. In cases where an enhancer overlaps a conserved block partially, the respective non-conserved boundary is interpolated by the distance to the closest conserved block. Accordingly, the size of the lifted enhancer region in mm9 will be approximately the same as the one of the respective mole enhancer. Nevertheless, to exclude artefacts, lifting is only accepted if the ratio of mole enhancer length / lifted length is <1.5. We define an enhancer sequence as conserved if the enhancer could be lifted successfully. In addition, we define an enhancer as conserved in enhancer function, if the mole enhancer overlaps a mouse enhancer irrespective of tissue-specificity.

### Transcriptomic analyses

For gene expression analysis, gonads from adult mice, P7 infants and embryos at E13.5, were dissected and RNA was extracted from these samples using the RNeasy Mini Kit (QIAGEN) according to the manufacturer’s instructions. For mole gonads previously published RNA-seq data was used^7^. The samples were sequenced using Illumina HiSeq technology according to standard procedures. Read mapping was performed with the STAR v2.6.1d software^43^. Read counts were created using the R function “summarizeOverlaps” and normalized to RPKM based on the number of uniquely mapped reads. For the analysis of differential expression between samples, the DESeq2 tool was used with default settings^44^.

### Definition of female testis part specific regions

In order to prioritize enhancers by their potential relevance in testis part tissue, we first ranked enhancer regions by the difference in enhancer probability (score in testis part vs mean of scores in testis + ovary part). We defined the putative target genes of each enhancer as the gene with the closest transcription start site to the center of the enhancer region within the same TAD. Based on the differential expression analysis (testis part vs testis + ovary part), each target gene is ranked by specific expression in ovotestis (log2 fold-change). Finally, enhancers are ranked jointly for functional importance in the female testis part by the mean rank of probability score and the rank of the putative target gene.

### HiC

Previously published datasets from mole embryonic limb buds and adult ovotestes were used to inspect the SALL1 regulatory domain^7^. Maps were visualized with the Juice box software.

Mouse HiC was obtained from publicly available high-resolution datasets from neuronal progenitor cells (NPCs)^17^. Maps were visualized with the Juice box software.

### 4C sequencing

Embryonic tissues were dissociated with trypsin, filtered through a cell strainer to obtain a single cell suspension and subsequently fixed in 2% formaldehyde. Mouse embryonic stem cells (mESCs) were detached from culture plates and fixed in the same way. Cells were counted and five million cells were snap-frozen and stored at −80°C until processing.

4C-seq libraries were prepared according to standard protocols^45^. For the initial digestion, NlaIII was used in *SALL1* experiments and BfaI was used in ITR-BAC ES cells. For the second digestion, DpnII was used for all experiments. A total of 1.6 mg of each library was amplified by PCR for each viewpoint with primers listed in **Supplementary Table 2.** The libraries were sequenced using Illumina HiSeq technology according to standard procedures. Raw reads were pre-processed and mapped to the reference genome (talOcc4) using BWA^46^. Finally, reads were summarized and normalized by coverage (RPM) for each fragment generated by neighboring restriction enzyme sites. The viewpoint and its flanking fragments (1.5 kb upstream and downstream) were removed for data visualization and a window of 10 fragments was used to smoothen the data.

The mouse virtual 4C profile was derived from a genome-wide HiC-map from NPCs^17^ by first extracting the intrachromosomal contact maps for the chromosomes of interest using Juicer tools v0.7.5^47^ (KR normalized, MAPQ>=30, 5kb resolution). Afterwards, only map entries with at least one bin overlapping the viewpoint (chr8:89,044,162 (*Sall1*) on mm10 were used for the virtual 4C profile.

### LACZ reporter assay in transgenic mice

LACZ reporter assays were conducted following the “Transgenic Mouse Assay” protocol from Vista Enhancer Browser^19^. Briefly, enhancer sequences were amplified by PCR from mole genomic DNA using primers listed in **Supplementary Table 2**. PCR products were cloned into the standard Gateway entry vector (pENTR/D-TOPO vector, Invitrogen) according to the manufacturer’s instructions. Clones were then transferred into the destination vector containing a Gateway cassette and a Hsp68 promoter coupled to a LacZ reporter gene. For microinjection into fertilized eggs, plasmid DNA was linearized with XhoI or HindIII and purified using Montage PCR filter units and Micropure EZ column (Millipore). For pronuclear injection of FVB embryos, DNA was diluted to a final concentration of 1.5-2 ng/μl and used in accordance with standard protocols approved by the Lawrence Berkeley National Laboratory. Embryos were harvested at embryonic day 13.5, dissected and fixed in 4% paraformaldehyde (PFA). Tissues were stained for 24 hours with freshly prepared staining solution, washed and post-fixed in 4% PFA.

### BAC transgenesis for overexpression of Sall1

*SALL1* coding sequence (CDS) was amplified from a vector containing the cDNA mouse sequence (Origen, cat. No. MC203471) with specific primers compatible with the attB gateway recombination system (Invitrogen). Through the gateway system, the generated product was introduced into a modified Wt1-BAC, containing piggyBac DNA transposon elements, as well as attL docking sites. The Wt1-BAC vector was kindly provided by Dr. Koopman and its further modification was performed according to their previously published method^20^. After introduction of the *SALL1* minigene, a eukaryotic antibiotic resistance (dual Neomycin-Kanamycin cassette) was introduced into the BAC vector through recombineering for transfection into ES cells according to the protocol previously described^48^. Primers are listed in **Supplementary Table 2.**

### BAC transfection into female ES cells

Blastocysts from C57BL/6J mice were used to derive mouse embryonic stem cells (mESCs) by growing them with culture medium supplemented with leukemia inhibitory factor (LIF), as well as FGF/Erk and Gsk3 pathway inhibitors (2i). The derived mESCs were genotyped for sex and a female line was expanded through co-culture with mouse embryonic fibroblasts (MEFs) for further experiments.

Female mESCs were co-transfected with 3 μg piggybac transposase and 500 ng of the modified Wt1-*SALL1*-piggyBac-Neo-BAC using Lipofectamine LTX (Invitrogen), as described in a previous publication^49^. After Geneticin-G418 selection (250 μg/ml) for 5 to 10 days, clones were picked and checked for successful BAC integration with 3 genotyping PCRs. A primer pair targeting each piggybac ITR (5’ITR and 3’ITR) was used as positive control, while a primer pair targeting the BAC vector was used as negative control to confirm integration mediated by transposition, instead of random insertion. Positive clones were expanded and additional genotyping was done by 4C-seq, to confirm genomic integrations site, as well as number of integrations, as described previously^45^.

### Gene ontology analyses

For Gene Ontology (GO) terms enrichment analysis PANTHER software was used^50^, selecting all the common upregulated genes for the testis part of the ovotestes and in the *Sall1*-overexpressing mouse mutants. A total of 56 genes were evaluated.

